# The microbiota and immune system non-genetically affect offspring phenotypes transgenerationally

**DOI:** 10.1101/2023.04.06.535940

**Authors:** Jordan C. Harris, Natalie A. Trigg, Bruktawit Goshu, Yuichi Yokoyama, Lenka Dohnalová, Ellen K. White, Adele Harman, Christoph A. Thaiss, Elizabeth A. Grice, Colin C. Conine, Taku Kambayashi

## Abstract

The host-microbiota relationship has evolved to shape mammalian processes, including immunity, metabolism, and development^1–3^. Host phenotypes change in direct response to microbial exposures by the individual. Here we show that the microbiota induces phenotypic change not only in the individual but also in their succeeding generations of progeny. We found that germ-free mice exhibit a robust sebum secretion defect and transcriptional changes in various organs, persisting across multiple generations despite microbial colonization and breeding with conventional mice. Host-microbe interactions could be involved in this process, since T cell-deficient mice, which display defective sebum secretion^4^, also transgenerationally transmit their phenotype to progeny. These phenotypes are inherited by progeny conceived during *in vitro* fertilization using germ-free sperm and eggs, demonstrating that epigenetic information in the gametes is required for phenotypic transmission. Accordingly, small non-coding RNAs that can regulate embryonic gene expression^5^ were strikingly and similarly altered in gametes of germ-free and T cell-deficient mice. Thus, we have uncovered a novel mechanism whereby the microbiota and immune system induce phenotypic changes in successive generations of offspring. This epigenetic form of inheritance could be advantageous for host adaptation to environmental perturbation, where phenotypic diversity can be introduced more rapidly than by genetic mutation.

## Main Text

Barrier sites including skin, gut and lung are responsible for responding to a wide variety of environmental perturbations, including exposure to pathogens, physical disruption, and altered nutrient homeostasis^6–11^. The ability of these tissues to adapt to changing environments is a key component of organismal viability. In the long term, natural selection and evolution allow for optimal adaptation to many of these environmental shifts, while more severe and abrupt changes, such as infection, garner more acute responses. Just as a stratified, keratinized layer of skin has evolved over long periods to provide a permanent external barrier, the presence of skin commensal bacteria and the mechanisms by which they prevent pathogenic invasion allows for a more short-term form of cutaneous defense^12,13^.

Phenotypic diversity induced by genetic mutations, which are randomly introduced and accrue slowly over time, may not efficiently allow acute adaptation to changing environmental conditions. In contrast, environmentally directed epigenetic gene regulation could serve as a more rapid adaptive mechanism to induce phenotypic changes. Organisms may “fine-tune” phenotypes in response to environmental factors, which in some cases can be transmitted to their offspring. Indeed, recent work in *C. elegans* demonstrated the transmission of environmentally dependent, persistent phenotypes across generations even in the absence of the initial environmental perturbation^14,15^. However, whether this phenomenon occurs in mammals has been controversial, though it has been recently established that mammalian phenotypes affected by parental diet can be transmitted to successive generations transgenerationally^5,16^. Yet, these transmitted phenotypes can be variable and subtle and thus, a robust and reliable readout to study these inheritance mechanisms will be invaluable in advancing the field by allowing for more mechanistic transgenerational studies.

We recently described a cutaneous immune-sebum circuit, whereby thymic stromal lymphopoietin (TSLP)-stimulated T cells can influence the ability of sebaceous glands to secrete sebum, an oily substance that promotes skin hydration, acidification, and anti-microbial defense^4,17,18^. Since T cells can be activated by microbial antigens and TSLP can be released from skin keratinocytes with stimulation by microbial products^2,19,20^, we hypothesized that the skin microbiota could trigger T cell activation and TSLP expression to induce sebum secretion, which would in turn control skin commensals. This feedback mechanism could maintain a delicately balanced skin ecosystem, which is essential for optimal barrier function^21,22^. In this study, we found that skin microbiota does indeed control sebum secretion, though not in an acute manner as originally hypothesized. Instead, we found that commensal microbes influence sebaceous gland function, as well as the transcriptional profiles in multiple organs by transgenerational epigenetic inheritance. Further, we find that T cells additionally regulate analogous inherited phenotypes transgenerationally, including defective sebum secretion. Both the microbiota and T cells strikingly influence the small regulatory RNA payload in gametes, generating modulated epigenetic information with the potential to transmit non-genetically inherited phenotypes. Our results reveal that the microbiome and immune system control epigenetic information in the gametes to modulate the phenotypes of succeeding generations of progeny.

## Results

### Reduced sebum secretion in germ-free mice

We have previously shown that germ free (GF) mice display abnormal epidermal structure and permeability barrier function^21,23^. To determine if the skin barrier defect carries to sebaceous gland (SG) function, we examined an RNA sequencing (RNA-seq) dataset generated by our lab (GSE162925) and found that GF epidermis showed reduced expression of lipid metabolism and anti-microbial peptide (AMP) genes, both processes important in SG biology (**fig. S1A**). Consistent with these findings, the amount of sebum present on the fur of GF mice was significantly reduced compared to conventionally-raised (CR) control mice (**Fig. 1A**). Moreover, RNA-seq of laser capture microdissection (LCM)-isolated SGs revealed a distinct transcriptional signature in GF mice (**Fig. 1B; fig. S1B**) with 45 genes significantly upregulated and 127 genes significantly downregulated in GF SGs (**Fig. 1C**). Gene ontology (GO) and gene set enrichment analysis (GSEA) showed that GF SGs displayed downregulation of lipid metabolism and cell death pathways, processes important in SG lipogenesis and holocrine (cell death-mediated) sebum secretion (**Fig. 1D; fig. S1C**)^24,25^.

**Fig. 1.**
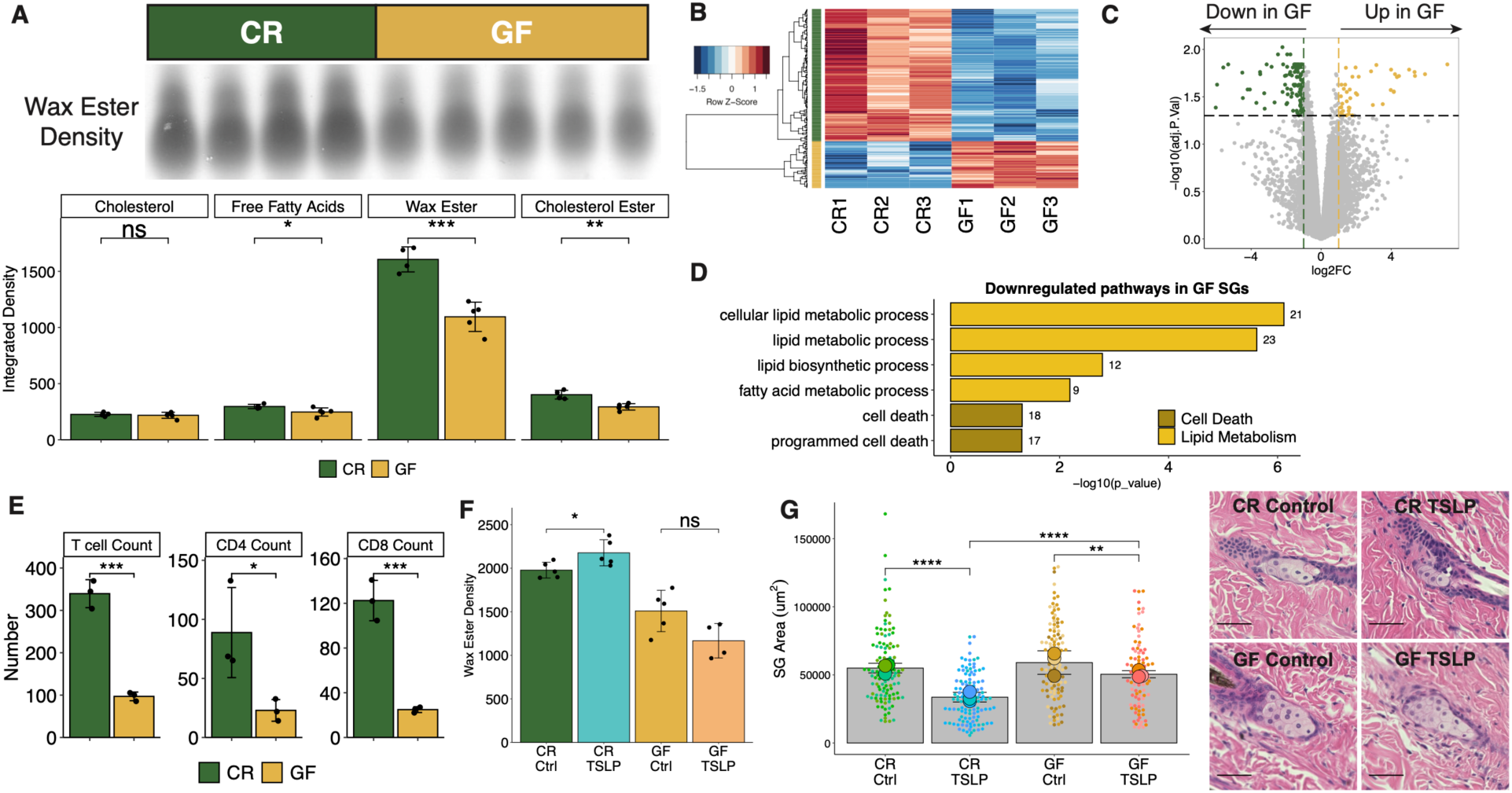
GF mice display a defective immune-sebum circuit. (**A**) Wax ester intensity and hair sebum lipid quantification by thin layer chromatography (TLC) (*n* = 4 or 5 mice/group). (**B**) Heatmap of DEGs in LCM-isolated GF or CR mouse SG. (**C**) Volcano plot highlighting SG genes upregulated (45 genes) and downregulated (127 genes) in GF mice. (**D**) Selected GO terms downregulated in GF SGs. Number of genes within the dataset within each term listed beside the bar. (**E**) Number of skin T cells (*n* = 3 mice/group). (**F** to **G**) CR or GF mice treated with AAV for 14 days. (F) TLC quantification of wax ester from hair (*n* = 4 or 5 mice/group). (G) SG area with representative H&E images (scale bars, 100 μm; *n* = 3 mice/group, *n* = 27 to 53 SGs/mouse). Sequencing experiments were performed once. All other experiments were performed 2-5 times. ns, not significant, \**p* < 0.05, \*\**p* < 0.01, \*\*\**p* < 0.001, \*\*\*\**p* < 0.0001 by Student *t* test. Data are shown as mean ± SD.

The cutaneous immune-sebum circuit is mediated by TSLP-stimulated T cells^4^. Accordingly, we found that GF skin exhibited significantly reduced T cell numbers, as well as a trend towards reduced TSLP expression compared to CR skin (**Fig. 1E**; **fig. S1D**). As we have previously shown that TSLP overexpression leads to sebum hypersecretion and SG size reduction (due to increased holocrine secretion)^4^, we tested whether TSLP overexpression could restore sebum secretion in GF mice. GF mice treated with TSLP showed unaltered sebum secretion and a less profound change in SG size (**Fig. 1F and G**). Together, these data suggest that GF mice harbor a defect in homeostatic sebum secretion that cannot be overcome by TSLP overexpression.

### Inheritance of defective sebum secretion

We next sought to correct the SG defect by microbial conventionalization of adult GF mice with bedding from CR mice (**fig. S2A**)^21^. After 8 weeks of colonization in adult mice, we found that sebum secretion remained defective (**Fig. 2A)**, despite adequate restoration of skin commensals (**fig. S2B**). Given that certain phenotypes are controlled by microbial exposure specifically during the neonatal period^1,26,27^, we tested whether microbial colonization in early life would restore sebum secretion (**fig. S2A**). Surprisingly, conventionalization of pregnant GF dames still gave rise to adult progeny with a sebum secretion defect (**Fig. 2B**). These data suggested that the sebum secretion defect may be inherited by the progeny of GF parents.

**Fig. 2.**
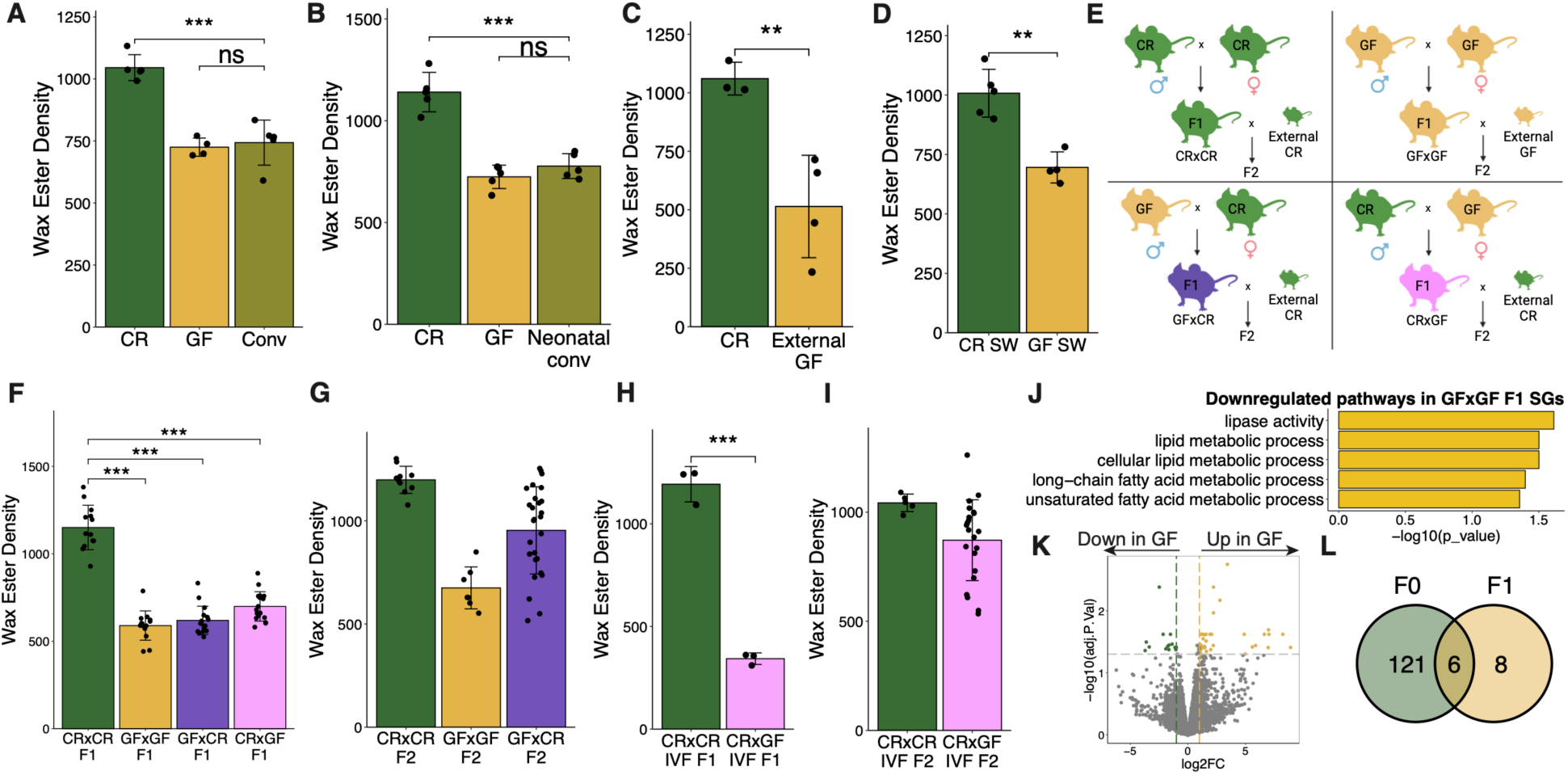
Sebum phenotypes are transmitted to progeny transgenerationally from GF mice. (**A** and **B**) TLC quantification of hair wax esters from CR, GF, and (A) 8 weeks post-conventionalized adult GF mice (*n* = 4 or 5 mice/group) or (B) GF mice conventionalized from birth (*n* = 4 or 5 mice/group). (**C** and **D**) TLC quantification of hair wax esters from (C) CR or GF mice from the UNC gnotobiotic core (*n* = 3 or 4 mice/group); (D) CR or GF Swiss-Webster mice (*n* = 4 or 5 mice/group) (**E**) Breeding scheme for transgenerational experiments. (**F** to **I**) TLC quantification of hair wax esters from combinatorial CR and GF natural breeding in the (F) F_1_ (*n* = 13 to 17 mice/group) and (G) F_2_ (*n* = 7 to 27 mice/group) generations and mice derived from IVF and the resulting (H) F_1_ (*n* = 3 mice/group) and (I) F_2_ (*n* = 6 or 22 mice/group) generations. (**J** to **L**) Gene expression by RNA-seq of F_1_ CR×CR and GF×GF SGs (*n* = 2 or 3 mice/group). (J) Selected GO terms downregulated in F_1_ GF×GF SGs. (K) Volcano plot highlighting DEGs in GF×GF F_1_ mice. (L) Downregulated genes shared between F_0_ and F_1_ GF SGs. Sequencing experiment performed once. All other experiments were performed 2-3 times. ns, not significant, \**p* < 0.05, \*\**p* < 0.01, \*\*\**p* < 0.001 by Student *t* test. Data are shown as mean ± SD.

We considered the possibility that GF mice (C57BL/6 strain) in our gnotobiotic facility (University of Pennsylvania) allopatrically acquired a genetic mutation that was responsible for preventing physiologic sebum secretion through genetic drift^28^. To test this, we examined sebum secretion in GF C57BL/6 mice from another gnotobiotic facility (University of North Carolina [UNC]) and in another GF strain (Swiss-Webster) from our facility. Both the GF C57BL/6 strain from UNC and the GF Swiss-Webster mice displayed a secretion sebum defect similar to GF C57BL/6 mice from our colony (**Fig. 2C** and **D**). Together, these data argued against a randomly acquired genetic mutation in GF mice as the cause of the sebum secretion defect.

To test if the sebum secretion defect was heritable, we bred CR mice with GF mice in a conventional animal facility in all four combinations: CR male × CR female (CR×CR), GF male × CR female (GF×CR), CR male × GF female (CR×GF), and GF male × GF female (GF×GF) (**Fig. 2E**). Despite similar microbial colonization (**fig. S2C**), the F_1_ progeny with at least one GF parent displayed a sebum secretion defect comparable to parental GF mice (**Fig. 2F**), suggesting that the GF sebum secretion phenotype is dominantly inherited. In some experiments, a small minority (16 of 148) of F_1_ mice with a GF parent displayed normal sebum secretion. To test if the phenotype persisted to the F_2_ generation, CR×CR and GF×CR F_1_ mice were bred to new CR mice while mice from the GF×GF group were bred to new GF mice as a negative control (**Fig. 2E**). Approximately half of the GF×CR group in the F_2_ generation remained defective, suggesting that the sebum secretion defect was transgenerationally inherited (**Fig. 2G**). Finally, to ensure that the phenotype can be transmitted by the gametes of GF mice in the absence of potentially confounding environmental factors, *in vitro* fertilization (IVF) with subsequent implantation into surrogate mothers was performed. Similar to results obtained with natural breeding, sebum secretion was defective in F_1_ progeny when eggs or sperm were of GF origin (**Fig. 2H**). In the F_2_ generation, the sebum secretion defect persisted in approximately one-third of F_2_ offspring of CR×GF IVF mice mated to CR mice (**Fig. 2I**).

To corroborate the persistent defective sebum secretion findings, we performed RNA-seq of LCM-isolated SGs from CR×CR and GF×GF F_1_ mice. Lipid metabolism-related pathways were downregulated in the GF×GF F_1_ SGs (**Fig. 2J**). Of the 14 downregulated genes, 6 were also downregulated in the SGs of parental GF mice (**Fig. 2K** and **L**). These results suggest that the sebum secretion phenotype is transmitted transgenerationally, and SG transcriptomic changes are transmitted at least intergenerationally, where the phenotype persists after removal of the environmental perturbation (in this case, the lack of microbiota) but is restored sporadically over time.

### Inherited phenotypes not limited to sebum secretion

To determine whether the transgenerational epigenetic inheritance process in GF mice extended to phenotypes other than sebum secretion, we first tested whether the immune-sebum circuit remained defective in progeny of GF mice. In the F_1_ generation, TSLP expression and T cell numbers continued to be reduced in the skin of offspring from colonized GF×GF mice (**fig. S2D and S2E**). Additionally, GF mice have a dysregulated skin transcriptome^21,23^ with downregulation of lipid-related genes and pathways^21^ (**fig. S1A**). Differentially expressed genes (DEGs) were also seen in the skin transcriptomic profile of GF×GF and GF×CR F_1_ mice compared to CR×CR F_1_ mice, suggesting that mice derived from a GF parent maintain altered cutaneous gene expression (**Fig. 3A**). 18 DEGs from the F_1_ generation persisted to the F_2_ generation in the GF×GF group (**Fig. 3B**). Some of these DEGs persisted, but lost significance in the GF×CR F_2_ mice (**Fig. 3B**) because the gene expression pattern in the F_2_ generation was bimodally distributed due to sporadic reversion of gene expression in a proportion of the progeny, mimicking the pattern of sebum inheritance. Two examples (*Erdr1* and *Hist1h4m*) are shown in the GF×CR F_2_ group, where the mice displayed a stochastic “on-or-off” level of expression (**Fig. 3C**).

**Fig. 3.**
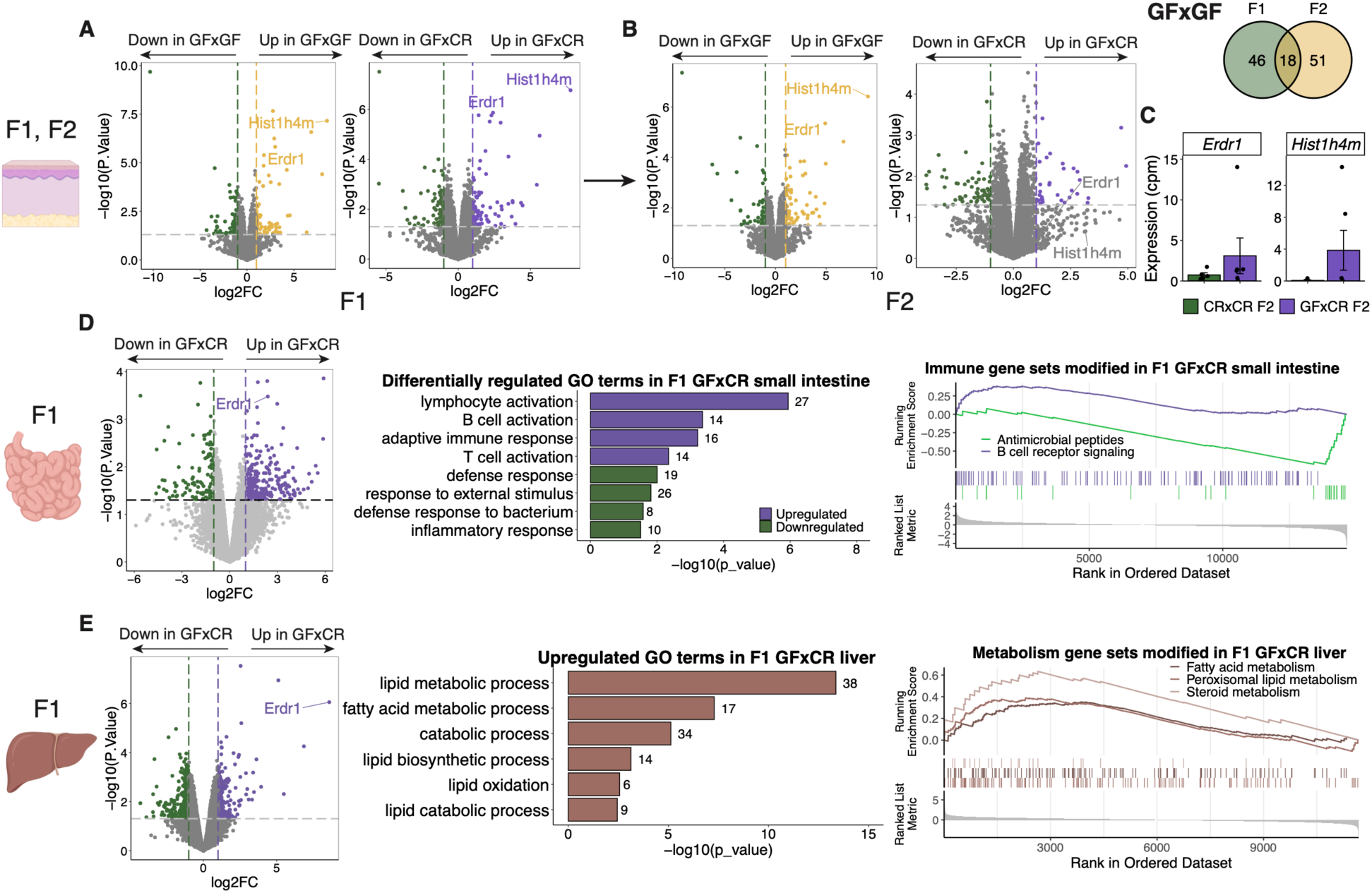
GF barrier and metabolic tissue display transgenerational transcriptional dysfunction. (**A** to **C**) Gene expression by RNA-seq of F_1_ and F_2_ CR×CR, GF×GF, and GF×CR back skin (*n* = 3 to 6 mice/group). Volcano plots representing pairwise group comparisons of DEGs across (A) F_1_ and (B) F_2_ generations, highlighting two common genes and a Venn diagram highlighting all common genes between F_1_ and F_2_ GF×GF skin. (C) Counts-per-million of two F_1_ DEGs with bimodal expression in F_2_. (**D**) Gene expression by RNA-seq of F_1_ CR×CR and GF×CR small intestine (*n* = 3 or 4 mice/group), including DEGs, GO terms, and GSEA showing upregulated and downregulated pathways. (**E**) Gene expression by RNA-seq of F_1_ and CR×CR and GF×CR liver tissue (*n* = 3 or 4 mice/group), including DEGs, GO terms, and GSEA showing upregulated pathways. The number of genes within the dataset within each term is listed beside the bar. Genes in GSEA plot shown in ranked order by running enrichment scores. Sequencing experiments were performed once. Data are shown as mean ± SD.

Next, we asked whether transcriptional changes are seen in tissues of GF×CR F_1_ progeny aside from skin. As proof-of-concept, we chose the small intestine, another barrier site, and the liver, a metabolic tissue, which have both been shown to be transcriptionally dysregulated in GF mice^3,29–32^. We performed a comparative transcriptomic analysis on CR×CR and GF×CR F_1_ tissues. In the small intestine of GF×CR F_1_ mice, immune activity related to innate bacterial defense pathways were downregulated, while adaptive and lymphocytic immune pathways were upregulated compared to CR×CR F_1_ mice (**Fig. 3D**), suggesting an alteration in immune response to microbes. In the liver, GF×CR F_1_ tissue displayed a change in metabolic function, with both lipid biosynthetic and catabolic processes upregulated, suggesting differential processing of lipid species in the GF×CR F_1_ mice (**Fig. 3E**). Taken together, these results suggest that the epigenetic inheritance pattern is not limited to SGs; multiple tissues in F_1_ mice derived from a GF parent are dysregulated even after colonization, suggesting that this process could represent a pervasive mechanism for controlling gene expression and phenotypes across generations of progeny.

### Small RNA profiling of GF gametes

To determine the mechanism by which GF parents can epigenetically alter the expression of genes and phenotypes in future generations, we took a step-wise approach to interrogate changes in GF gametes and F_1_ embryos. We hypothesized that the microbiota could have an impact on gametes and embryos, informing the developing offspring of the microbial environmental status. It has recently been discovered that changes in small non-coding RNA (ncRNA) in sperm can lead to alterations in embryonic gene regulation and phenotypes of future generations^5,16, 33–35^. In particular, the tRNA fragments (tRFs) Gly-GCC, Val-CAC and a subset of miRNAs have been shown to be delivered to sperm by fusion with extracellular vesicles, called epididymosomes^5^. Further, tRF-Gly-GCC and epididymally-acquired miRNAs have been demonstrated to regulate embryonic gene expression post-fertilization as well as to program offspring phenotypes^16,36^. To this end, we examined the small ncRNA profile of sperm and eggs of CR and GF mice using small RNA-sequencing. We found a significant increase in many miRNAs and a decrease in specific tRFs in GF compared to CR sperm, including decreased expression of tRF-Val-CAC (**Fig. 4A; fig. S3A**). Similarly, in GF eggs, we most notably observed a widespread reduction in tRFs (**Fig. 4B**), including 21/24 tRFs which are also downregulated in GF sperm and 2 shared upregulated miRNAs (**Fig. 4C**). To note, the sperm and egg small RNAs were sequenced using distinct protocols, as only small quantities of eggs can be retrieved, thus a protocol optimized for low amounts of input RNA was used for egg RNA sequencing. Nevertheless, the finding that common small RNAs are regulated in the sperm and eggs of GF mice, and their shared transmitted phenotypes, is striking. These data suggest that there may be shared small ncRNA differences in male and female gametes that lead to changes in inheritable phenotypes of GF mice.

**Fig. 4.**
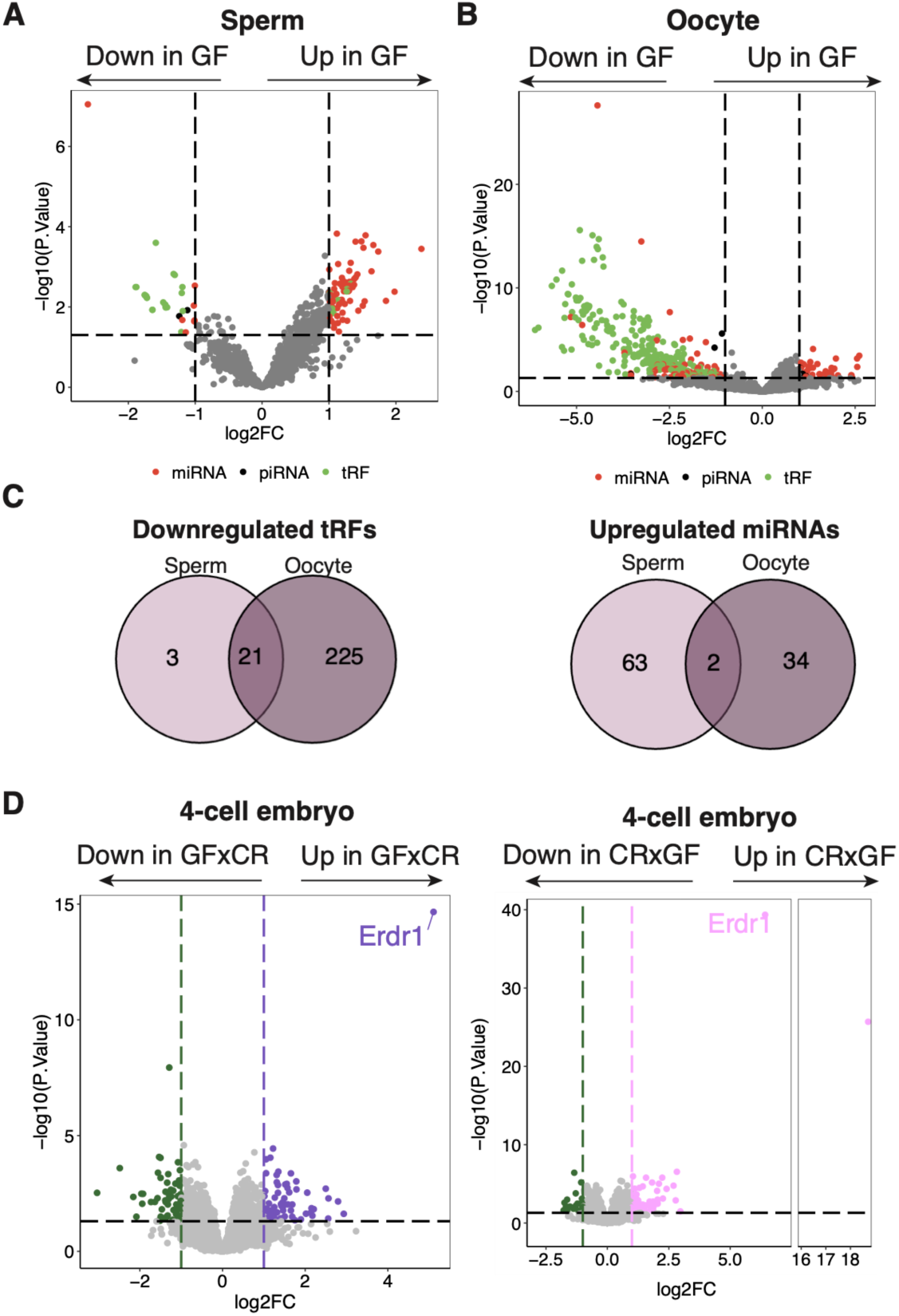
Sperm and eggs from GF mice exhibit a distinct small RNA profile that correlates with changes in early embryonic gene expression. (**A**) Small RNA expression by RNA-seq of CR and GF sperm (n = 8 mice/group), highlighting 72 upregulated and 33 downregulated RNAs in GF sperm. (**B**) Small RNA expression by RNA-seq of CR and GF eggs (n = 3 replicates of ∼20 eggs), highlighting 311 downregulated and 39 upregulated RNAs in GF eggs (**C**) Downregulated tRFs and upregulated miRNAs shared by GF sperm and eggs. (**D**) Gene expression by RNA-seq of CR×CR vs GF×CR, and CR×CR vs CR×GF 4-cell embryos (*n* = 9 to 25 embryos/group, collected over three biological replicates of IVF). DEGs defined as log_2_-fold change > 1 or < −1, *p* value < 0.05. All sequencing experiments were performed once.

We next determined whether fertilization with GF gametes caused altered gene expression in the early embryo. Single embryo mRNA-sequencing was performed at the 4-cell stage after IVF of CR or GF eggs with either GF or CR sperm and morula stage after IVF of CR eggs with either GF or CR sperm. We found significant transcriptional changes in embryos at both stages, with 79 upregulated and 48 downregulated genes in GF×CR 4-cell embryos, and 223 upregulated and 179 downregulated genes in CR×GF 4-cell embryos compared to CR×CR 4-cell embryos (**Fig. 4D; fig. S3B**). Of these genes, notably significant was *Erdr1*, which was also seen as a commonly dysregulated gene in adult somatic tissues (**Fig. 3**). The function of *Erdr1* is thought to be related to regulation of cell death, proliferation and migration, thus, extreme changes in *Erdr1* expression could lead to significant embryologic changes^37–39^.

### Role of T cells in epigenetic inheritance

Similar to GF mice, we have previously reported that homeostatic sebum secretion is reduced in *Rag2^−/−^* (which lack T and B cells) and *TCRβ^−/−^* (which lack αβ T cells) mice^4^. To test whether this defect was acutely restorable, we adoptively transferred T cells into *Rag2^−/−^* mice. Similar to colonization of GF mice, homeostatic sebum secretion was not restored (**Fig. 5A**). To see if the sebum secretion defect was transmissible to progeny, we crossed *Rag2^−/−^* males or females to wildtype (WT) females or males to create F_1_ *Rag2^+/−^* mice, which have a normal T cell compartment^40^. Similar to GF mice crossed to CR mice, we found that sebum secretion was defective in F_1_ *Rag2^+/−^* mice (33 of 36 mice across 4 experiments) (**Fig. 5B; fig S4**). We next crossed the F_1_ *Rag2^+/−^* mice against each other to generate progeny of all 3 genotypes (*Rag2^−/−^*, *Rag2^+/−^*, and *Rag2^+/+^*). There was a mix of F_2_ progeny with either normal or defective sebum secretion, regardless of genotype; a fraction of genotypically WT mice showed defective sebum secretion, while *Rag2^−/−^* mice showed normal sebum secretion, indicating that the phenotype of sebum secretion is not correlated with genotypes, but rather with parental immune status (**Fig. 5C**). Similar to *Rag2^−/−^* mice, the F_1_ progeny of *TCRβ^−/−^* mice crossed to WT mice also showed defective sebum secretion (**Fig. 5D**), suggesting that the absence of T cells was responsible for transgenerational epigenetic inheritance of the sebum secretion defect.

**Fig. 5.**
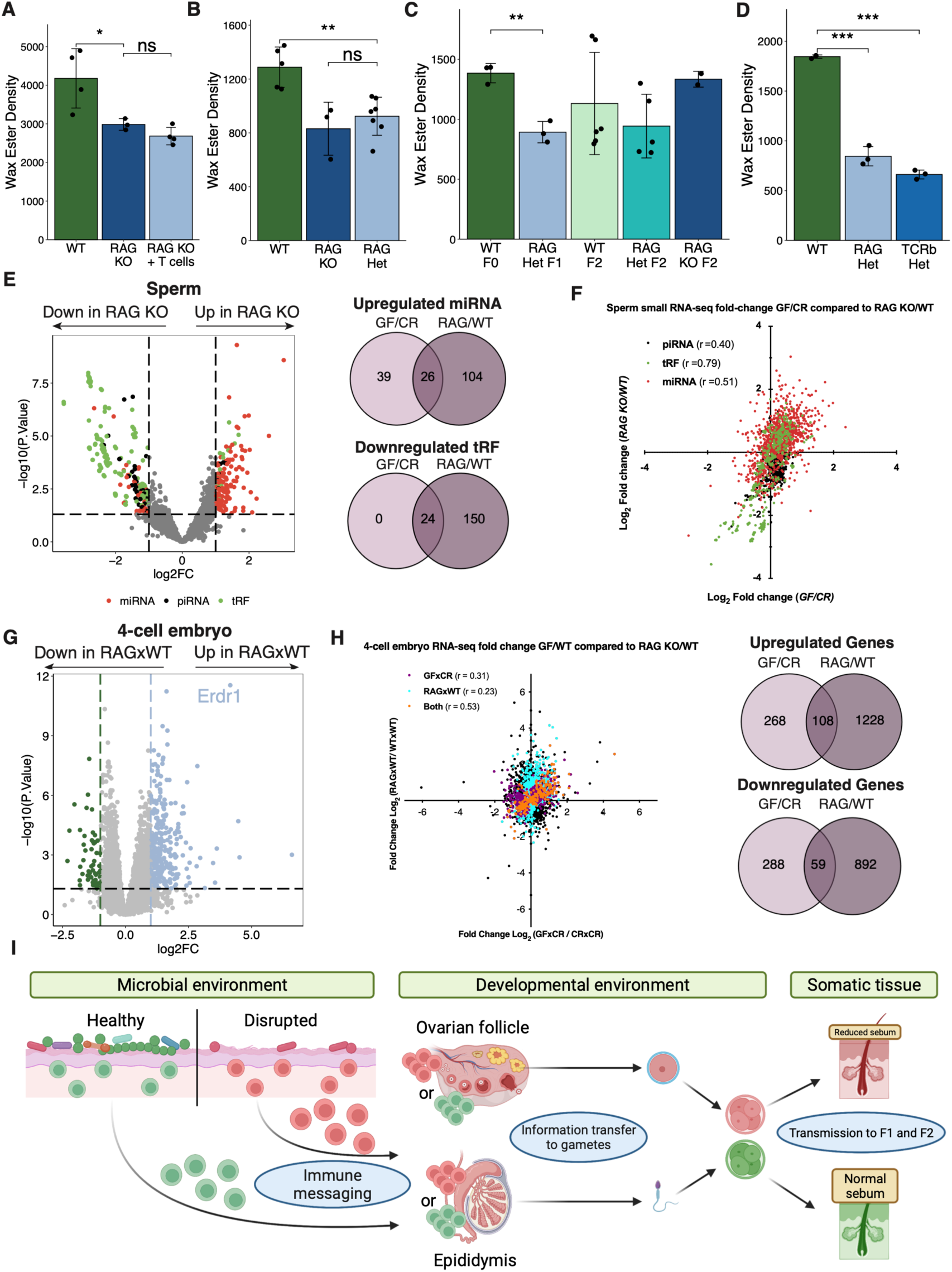
T cell deficient mice display a dysregulated sperm small RNA profile and a sebum secretion defect that is transmitted to progeny transgenerationally. (**A** to **D**) TLC quantification of hair wax esters from (A) WT or *Rag2^−/−^* mice with or without reconstitution by T cells (*n* = 3 or 4 mice/group). (B) WT, *Rag2^−/−^*, and F_1_ *Rag2^+/−^* mice (*n* = 3 to 7 mice/group). (C) WT, F_1_ *Rag2^+/−^*, and F_2_ WT, *Rag2^+/−^*, and *Rag2^−/−^* mice (*n* = 2 to 6 mice/group). (D) F_1_ WT, *Rag2^+/−^*, and *TCRβ^+/−^* mice (*n* = 3 mice/group). (**E**) Small RNA expression by RNA-seq of WT and *Rag2^−/−^* sperm (*n* = 6 or 8 mice/group), highlighting 149 upregulated and 235 downregulated RNAs in *Rag2^-/-^*sperm with upregulated miRNA and downregulated tRFs shared between GF and *Rag2^−/−^* sperm quantified. (**F**) Pearson correlation analysis of all GF vs *Rag2^−/−^* sperm small RNA differential abundance. (**G**) Gene expression by RNA-seq of WT and RAG×WT 4-cell embryos (*n* = 21 or 25 embryos/group, collected over three biological replicates of IVF). (**H**) Pearson correlation analysis of GF vs *Rag2^−/−^* embryo gene expression on genes filtered for *p* value < 0.05 (purple: significant only in GF derived embryos; blue: significant only in *Rag2^−/−^* derived embryos; orange: significant in both GF and *Rag2^−/−^* derived embryos), with DEGs shared between GF and *Rag2^−/−^* embryos quantified. (**I**) Model for microbial and immune-mediated transgenerational epigenetic inheritance. Sequencing experiments performed once. All other experiments were performed 2 or 3 times. ns, not significant, \**p* < 0.05, \*\**p* < 0.01, \*\*\**p* < 0.001 by Student *t* test. Data are shown as mean ± SD.

We next profiled the small ncRNA profile of sperm of *Rag2^−/−^* mice. Like GF mice, profound changes in small ncRNAs levels were identified (**Fig. 5E; fig. S3A**). There was a high degree of concordance as shown by the *r* value between the small RNA log_2_-fold change in the sperm of GF and *Rag2^−/−^* mice in all small RNAs compared to their respective controls, with all tRFs downregulated in GF mice also found downregulated in *Rag2^−/−^* mice (**Fig. 5E** and **F**). To determine whether *Rag2^−/−^* embryos follow a similar concordance, we performed single embryo RNA-seq on embryos generated by *Rag2^−/−^* sperm and WT eggs (RAG×WT). We observed many differentially expressed genes in *Rag2^−/−^* derived 4-cell- and morula-stage embryos with 167 significantly changed genes shared between *Rag2^−/−^* and GF derived 4-cell embryos (**Fig. 5G; fig. S3C**). However, there were both commonly shared and distinct DEGs identified between GF and *Rag2^−/−^* 4-cell embryos, suggesting there may be both interdependent and independent contributions of the microbiota and the adaptive immune system in controlling paternal epigenetic inheritance patterns (**Fig. 5H**).

## Discussion

The results presented here describe a microbial and immune-dependent form of transgenerational epigenetic inheritance with the ability to influence the phenotypic diversity of future generations. Our data provide evidence that the commensal microbiota is not only important for acute changes in organ function but can have a persistent effect on future generations. We also describe a new and impactful role of the immune system in influencing gametes to alter the control of gene expression and phenotypes of succeeding generations of progeny.

There are many examples illustrating the importance of host-microbe interactions in regulating functional biological processes^21,26,27^. As such, there are innumerable defects present in the tissues of GF mice, ranging from barrier sites to internal organ systems^3,21,41^. Reversal of these GF defects with bacterial colonization is a common experimental tool in microbiome research, though some GF phenotypes are not acutely reversible with colonization, without explanation. From the work we describe, we propose this dichotomy in the reversal of GF phenotypes is due to an important distinction in acute versus persistent epigenetically inherited phenotypes. As we have seen striking evidence that a dysregulated GF transcriptome in multiple tissues is passed across generations, it will be important to match the transcriptional changes to phenotypic function of these tissues to determine their effects. For example, the gut immune function of GF×CR F_1_ mice is likely to be perturbed, given the reduction in transcriptional programs that control innate bacterial response in the small intestine of conventionalized GF-derived F_1_ mice.

While we hypothesized that microbes use T cells as messengers to communicate with reproductive tissues, we did not observe a perfect correlation between the gene expression changes in progeny resulting from the lack of microbiota and T cells, suggesting that there are independent effects of both systems in controlling transgenerational phenotypes. Yet, based on many similarities in the inheritance pattern, analogous small RNA changes in the gametes, and related inherited modulation of embryonic gene expression, we propose a model (**Fig. 5I**) whereby the microbial environment at least in part is detected by T cells. T cells can then transfer this information to gametes, altering the epigenetic information in the form of small ncRNAs, transmitting phenotypic diversity to future generations.

It has been shown that the microbial composition of barrier sites is altered in response to the environment^42,43^. Thus, environmental perturbations could be sensed by changes in commensal microbial composition, which then is provided as information to offspring to adapt more successfully to the environment. Moreover, previous reports in mice have described how diet alterations and stress program non-genetically inherited phenotypes in subsequent generations of progeny^5,16,44^. The microbiome and the immune system have been independently linked to both changes in diet and stress^45–47^ and thus it is plausible that modifications in diet or introduction of persistent stress and the resulting microbial and immune alterations are responsible for altering epigenetic information in the gametes and intergenerational information transfer. Teleologically, we believe our observations suggest microbial presence can provide environmental context to offspring to allow for optimal use of energy and metabolism. As an example, we show that F_0_ and F_1_ GF mice have reduced sebum secretion, and that F_1_ livers show altered metabolic lipid processing. It is enticing to speculate that because of the absence of microbes in GF mice, the host is shunting metabolic effort normally reserved for sebum secretion and barrier function to the liver to save energy.

Despite numerous examples in model organisms ranging from plants to *C. elegans*^48^, the existence of transgenerational epigenetic inheritance in mammals has been controversial. This controversy is a result of weakly penetrant and expressive phenotypes that have been demonstrated to be transmitted by transgenerational epigenetic inheritance and undefined molecular mechanisms underlying the phenomenon. Only recently has evidence of this pattern of inheritance contributing to mammalian phenotypes been uncovered, yet these studies have not completely resolved the controversy^14,49–58^. Sperm RNAs have been demonstrated to transmit epigenetically inherited information to offspring in a variety of organisms, including mammals, in an assortment of intergenerational inheritance paradigms^15,33, 59–63^. However, because the study of gametic RNAs programming phenotypes in the progeny is still in its infancy, the mechanism by which these RNAs can produce changes to development that manifest as non-genetically inherited phenotypes is poorly understood. We describe a robust readout of transgenerational epigenetic inheritance patterns using sebum secretion, providing a sensitive model for groundbreaking studies to understand the molecular mechanisms underlying the transfer of epigenetic information between generations and throughout development.

From our studies, we observe that the gene *Erdr1* is strikingly upregulated in both GF-derived early embryos and adult somatic tissues. Interestingly we also find that *Erdr1* is regulated analogously in early embryos derived from *Rag2^−/−^* crossed with WT female mice. While the significance of these observations is currently unknown, *Erdr1* poses as an intriguing target for future studies of microbial-dependent transgenerational phenotypes, potentially acting as a common thread across generations.

As a result of the discovery of this microbial-immune-transgenerational phenotypic inheritance, we could contextualize events from the past, attempt to explain the state of human health in the present, and learn how our current decisions could affect the future. In the modern era, one of the most significant changes in human health is the explosive onset of atopic and autoimmune disease. The “hygiene hypothesis” is a popular idea to explain how the prevalence of atopy and autoimmunity have risen, whereby human society has become more hygienic and less barraged by pathogens to train the immune system, leading to immune overactivation in the form of allergy and autoimmunity^64,65^. We might consider that the effects of sanitation from industrialization have been passed down over multiple generations and are increasingly materializing in the modern day in the form of immune dysregulation. As a form of positive adaptation, this phenomenon may be a way for mammals to introduce phenotypic diversity into their offspring due to environmental change without the long-term necessity of genetic mutation-based natural selection. In this way, animals would have an increased chance at quickly and persistently adapting to new environmental threats looming on the horizon.

## Methods

### Animal models

All specific pathogen-free (conventionally raised: CR) mice used in these studies were derived from C57BL/6 mice purchased from Charles River Laboratories (strain number 556) unless otherwise specified. Germ-free (GF) mice were obtained from the University of Pennsylvania Gnotobiotic Core, which houses C57BL/6 and Swiss-Webster colonies in sterile isolators. Additional GF mice were obtained from the Gnotobiotic Core at University of North Carolina at Chapel Hill for comparing colony phenotypes. CR and GF breeding pairs were established in a conventional mouse facility at the University of Pennsylvania. *Rag2^−/−^* mice (Jackson Laboratories strain number 008449) and *TCRβ^−/−^* mice (Jackson Laboratories strain number 002116) were obtained from Jackson Laboratories and bred within our mouse facility. *Rag2^+/−^* and *TCRβ^+/−^* mice were derived in our animal facility by breeding the knockout strains to C57BL/6 wild-type mice (Charles River strain number 556). Unless otherwise specified, all mice used in these studies were 8 weeks old at the time of use. A combination of both male and female mice was used in the studies to ensure conclusions could be generalized to both sexes. All mice were housed in either specific pathogen-free or germ-free conditions and were handled under strict compliance with the University of Pennsylvania Institutional Animal Care and Use Committee regulations.

### Lipid extraction and thin layer chromatography

To isolate sebum lipids from mouse fur, a standardized 3 cm × 3 cm area of fur was shaved from the back. Fur was submerged in 2 mL of 2:1 (v/v) chloroform:methanol (Sigma 288306 and Sigma 322415) followed by sonication in a water bath for six minutes to dislodge lipids, and syringe filtration to remove fur from solution. Fur was then submerged in 2 mL of acetone and sonication and filtration steps were repeated. The organic solution containing fur lipids was evaporated using nitrogen gas until completely dry and dissolved in 250 μL of 4:1 chloroform:methanol (v/v). 5 μL of lipid solution was then loaded onto a thin layer chromatography plate (Sigma, 100390) and placed sequentially in (1) a shallow solution of 80:20:1 hexane (Sigma 296090):isopropyl diether (Sigma 673803):acetic acid to migrate to a plate height of 50%, (2) a shallow solution of 1:1 hexane:benzene (Sigma 401765) to migrate to a plate height of 80%, and (3) a shallow solution of hexane to migrate to a plate height of 90%, with 15 minutes of drying time between each migration step. Plates were then uniformly coated with 10% cupric sulfate (Sigma 451657)/8% phosphoric acid (Sigma P6550) solution, allowed to dry and baked at 120°C for 20 minutes to visualize lipid species. Adobe Photoshop was used to quantify the integrated density of the lipid bands. As a standard for lipid species identification, the TLC non-polar lipid mixture A (Matreya 1129) was used.

### Skin RNA extraction and cDNA synthesis for qPCR

On the day of tissue harvest, fur was shaved, back skin was removed, minced, snap frozen and stored at −80°C until further processing. To isolate RNA from skin, frozen tissue was transferred to TissueTube TT05M XT tissue bags (Covaris, 520140) and pulverized using a Covaris automated dry pulverizer (Covaris CP02) by submerging the tissue bag in liquid nitrogen for 10 seconds and immediately transferring for pulverization. Pulverized tissue was then transferred to 1 mL of TRIzol (ThermoFisher 15596026) and RNA extracted according to the TRIzol manufacturer’s protocols. Glycogen (Invitrogen 15-596-018) was used as a carrier during extraction. A Nanodrop 1000 was used to quantify isolated RNA. Following RNA extraction, cDNA was synthesized using Superscript Vilo (ThermoFisher 11754050) according to the manufacturer’s instructions. Quantitative polymerase chain reaction (qPCR) was then performed using the Taqman Fast Advanced Master Mix (Taqman 4444557) according to the manufacturer’s instructions, with the following primer from Taqman: *Tslp* (Mm01157588_m1). qPCR reactions were performed using a ViiA7 Real-Time PCR instrument (ThermoFisher).

### Flow cytometry

To quantify T cells in ear skin, dermal sheets were separated, and finely minced in RPMI 1640 media (ThermoFisher 11875093) complemented with 10% fetal bovine serum (R&D Systems S11150) (cRPMI) containing 100 μg/mL of Liberase TL (Roche 5401020001) and 50 μg/mL of DNase I (Sigma DN25). Minced tissue was incubated with shaking at 37°C for 1 hour and then strained through a 70 μm filter into a new tube containing 1 mL cRPMI. Cells were stained with cell surface stains and live-dead stain at 4°C for 15 minutes in PBS. Flow cytometry was then performed using an LSR II or LSR Fortessa instrument (BD Biosciences). Compensation was performed using compensation beads (BD Biosciences 552845). Flow cytometry data was analyzed using FlowJo software (TreeStar). Staining antibodies used included CD45.2 (mouse, PE fluorochrome, clone 104, BD Biosciences 560695, 1:200 dilution), TCRβ (mouse, PE-Cy7 fluorochrome, clone H57-597, BioLegend 109222, 1:200 dilution), CD4 (mouse, FITC fluorochrome, clone RM4-5, BioLegend 100510, 1:200 dilution), CD8a (mouse, PerCP-Cy5.5 fluorochrome, clone 53-6.7, BioLegend 100734, 1:200 dilution) and Live/Dead Near-IR (ThermoFisher L10119, 1:1000 dilution). CountBright beads were used for counting cells and normalization (ThermoFisher C36950).

### Laser capture microdissection, RNA extraction, and sequencing

Mouse back skin from CR, GF, CR×CR F_1_, and GF×GF F_1_ was collected and fixed overnight in 4% paraformaldehyde (Fisher AAJ19943K2) at 4°C followed by paraffin embedding. Laser capture microdissection (LCM) was performed using the LMD 7000 system (Leica Microsystems Inc., Chicago, IL). FFPE mouse skin was processed and cut onto a polyethylene naphthalate (PEN) slide designed for LCM processing (Leica 11505158). At least 1,000 SGs or 1,000,000 μm^2^ of tissue was isolated to obtain enough material for RNA extraction. SG RNA was extracted from post-LCM tissue using a Qiagen All Prep DNA/RNA FFPE Kit (Qiagen 80234). RNA concentration was measured by Qubit fluorometric quantification (ThermoFisher Qubit 2.0 Fluorometer) and RNA quality measured via BioAnalyzer (Agilent 2100 Bioanalyzer Instrument). cDNA libraries were prepared using Illumina Stranded Total RNA Prep with Ribo-Zero Plus Kit (Illumina 20040529) with IDT for Illumina RNA UD Indexes, Set A (Illumina 20040553). Libraries were assessed for cDNA quantity and library quality using Qubit and BioAnalyzer. As necessary, an extra bead wash step was performed to remove excess primer dimers in the library and purify samples further. Samples were then pooled and sequenced on a Nextseq 550 using a NextSeq 500/550 High Output Kit v2.5 (150 Cycles) (Illumina 20024907).

### Somatic tissue RNA-seq analysis

Transcriptomic analysis of sebaceous glands, skin, small intestine, and liver was performed in the R statistical computing environment version 4.2 and RStudio version 2022.02.1 using a pipeline adopted from an open-source toolkit for RNA sequencing analysis^66^. For pseudoalignment of reads to a reference genome, Kallisto was used in combination with the Ensembl species-specific database for gene annotation^67^. A filtration cutoff was used of 1 count per million in the number of samples equal to the *n* of the smallest group. Data was normalized using the Trimmed Mean of M-values (TMM) method from the EdgeR package^68^. Post-filtered, post-normalized data was then variance stabilized using the VOOM function from the Limma package^69^. Limma was then used for differential gene expression (DGE) testing with multiple testing correction via the Benjamini-Hochberg method^70^. For F_1_ SG samples, DGEs were defined as genes with BH-adjusted *p*-value < 0.05. For other somatic F_1_ and F_2_ samples a less stringent cut-off was used to define DGEs as genes with *p*-value < 0.05, as we were testing for broad similarities between cross-generational transcriptomic profiles. Gene ontology (GO) analysis was performed using the gprofiler2 R package with terms identified from the GO knowledgebase with FDR adj-p-value < 0.05 and gene set enrichment analysis was performed using the msigdbr, clusterprofiler, and enrichplot R packages^71–73^. For plots with points of extreme log_2_FC or significance, in order to visualize the data appropriately, the package R ggbreak was used^74^.

### Bacterial colonization and culture

To colonize germ-free mice with a conventional microbiota, germ-free mice were transferred to a conventional specific pathogen-free mouse facility and were exposed to bedding and cage material from three other mature mouse cages three times in the first week of transfer. The conventionalized germ-free mice had weekly cage changes, thus allowing for further microbial exposure. These mice were housed in this manner for eight weeks prior to takedown at which point mice were swabbed for bacterial culture and confirmation of adequate colonization. Swabs were dipped in PBS before deeply swabbing pre-shaved mouse back fur 10-15 times. Swabs were stored in PBS at RT for 30 minutes and then serially diluted for plating on blood agar. CFUs quantified by counting number of colonies on blood agar at a dilution with colony number between 10-100 and calculated based on dilution and volume used for plating.

### TSLP-AAV injections

Two adeno-associated virus vectors used in these studies were generated by the Penn Vector Core, including Control-AAV (AAV8.TBG.PI.eGFP.WPRE.bGH) and TSLP-AAV (AAV8.TBG.PI.mTSLP.IRES.eGFP.WPRE.bGH). Doses were previously optimized and mice were intravenously injected with 5×10^10^ genome copies of both Control- and TSLP-AAV for 14 days with serum TSLP levels confirmed using a murine specific ELISA (R&D Systems MTLP00)^4^.

### Histology

Skin tissue was isolated from mouse back and fixed at 4°C overnight in 4% paraformaldehyde (Fisher AAJ19943K2) prior to paraffin embedding. Processing and staining (H&E) was performed by the University of Pennsylvania’s Skin Biology and Disease Resource Center. For H&E skin sections, full section stitching at 40× magnification was performed to image one full section of skin per biological replicate. Within these sections, a total of 27 to 53 SGs were measured. For measurement of SG area, ImageJ (NIH) was used to draw circumscribing ellipses around SG edges. Samples were imaged using a Keyence VHX-6000 digital microscope system and were prepared for publication using ImageJ and Photoshop.

### Adoptive transfers

T cell adoptive transfers were performed by isolating CR or GF T cells for intravenous injection into *Rag2^−/−^* mice. Splenic T cells were isolated for transfer using a T cell negative selection kit (STEMCELL Technologies 19851) according to the manufacturer’s instructions. Intravenous injection of 5×10^6^ isolated T cells was then performed and six weeks later sebum secretion was measured.

### Bulk RNA extraction and sequencing

#### Skin

On the day of tissue harvest, fur was shaved, back skin was removed, minced, snap frozen and stored at −80°C until further processing. RNA was extracted from skin using the same method as described above in preparation for qPCR. *Small intestine*: On the day of tissue harvest, 1 cm of distal ileum was snap frozen and stored at −80°C until further processing. Tissue was homogenized in tubes with metal beads using TRIzol extraction as detailed previously.

#### Liver

On the day of tissue harvest, liver tissue was snap frozen and stored at −80°C until further processing. RNA was extracted from tissue using TRIzol extraction as detailed previously.

Quality control of RNA was performed using a Qubit fluorometer (ThermoFisher Qubit 2.0 Fluorometer) for quantification and Bioanalyzer (Agilent 2100 Bioanalyzer Instrument) or TapeStation (Agilent 4200) for RNA quality. *Skin*: cDNA libraries were prepared using the Illumina Stranded Total RNA Prep with Ribo-Zero Plus Kit (Illumina 20040529) with IDT for Illumina RNA UD Indexes, Set A (Illumina 20040553) and sequenced on a NovaSeq 6000 using a NovaSeq 6000 SP Reagent Kit v1.5 (100 cycles) (Illumina 20028401). *Small intestine*: Libraries were prepared using the Illumina Stranded mRNA Prep, Ligation kit according to the manufacturer’s instructions. Unique Illumina TruSeq dual indices were used for sample identification. Library pool was sequenced on an Illumina NextSeq 550 instrument using 75 cycles, single-end. *Liver*: mRNA-sequencing libraries were generated using Illumina stranded mRNA kit as per manufacturer’s instructions. Paired-end sequencing was performed using Illumina NextSeq 1000. Data were mapped using RSEM and normalized using transcripts per million (tpm).

### Egg collection*, in vitro* fertilization, and embryo culture and transfer

Eggs were retrieved from the ampullae of 4- to 6-week-old female mice following superovulation as previously described^75^. For egg collection for small RNA sequencing, cumulus oocyte complexes (COCs) were incubated in hyaluronidase (1 mg/ml) to dissociate cumulus cells from eggs. Eggs were washed through six droplets of KSOM to remove any residual cumulus cells and collected in 1× TCL buffer (supplemented with 1% β-mercaptoethanol). For *in vitro* fertilization (IVF), spermatozoa were collected and capacitated as previously described^75^. Spermatozoa (2 × 10^5^) were added to the IVF droplet and co-incubated with eggs for 3 h at 37°C under an atmosphere of 5% O_2_, 6% CO_2_. Presumptive zygotes were washed in KSOM and cultured until 2-cell (24 h), 4-cell (46 h) and morula (72 h) stage.

Transfer of IVF generated embryos was performed at the Children’s Hospital of Philadelphia Transgenic Core. Embryos cultured to the 2-cell stage of development were transferred to the oviduct of pseudopregnant recipient females to produce live pups.

### Cauda sperm isolation and RNA extraction

Mature populations of cauda spermatozoa were isolated from adult (8-12-week-old) mice and total RNA was extracted as previously described^5^. In brief, sperm pellets were resuspended in lysis buffer supplemented with 33 mM DTT and 0.66 mg/mL Proteinase K and incubated at 60°C for 15 min. After incubation, an equal volume of water and TRIzol was added to each sample. The samples were transferred to phase lock tubes for phase separation and precipitation as above. Small RNA was purified using denaturing polyacrylamide gel electrophoresis (PAGE)^5^.

### Small-RNA sequencing

The sperm small RNA profile was cloned using the Small RNA TruSeq kit (Illumina). Adapter ligation and conversion to DNA was performed as per the manufacturer’s instructions. Amplification of libraries was optimized for representative samples to determine the optimum number of cycles. Once cycle number was determined all libraires were amplified and gel-purified. Libraries were combined at equimolar concentrations and sequenced using Illumina NextSeq 1000 (75 cycles). For small RNA sequencing of eggs, three replicates of ∼ 20 eggs were collected and lysed in 1 × TCL buffer. After RNA extraction (TRIzol, phase separation and precipitation), RNA was DNase treated and reconstituted in nuclease free water. Total RNA was then used for library preparation using the SMARTer small RNA (smRNA)-seq kit for Illumina (Takara Bio) according to the manufacturer’s instructions. After library preparation and bead clean up, individual samples were combined at equimolar concentrations and purified using non-denaturing PAGE to purify cloned, adapter-ligated small RNA constructs. Finally, pooled libraries were sequenced using Illumina NextSeq 1000 (75 cycles). Raw read count data was normalized and analyzed for differentially abundant small RNAs as previously outlined^5^.

### Embryo mRNA-sequencing

Single embryos, sired by control, germ-free or *Rag2*^−*/*−^ sperm or eggs from control or GF mice were collected for single-embryo/egg RNA-sequencing. Embryos were transferred to a 96-well plate and fresh 1 × TCL buffer with 1% β-mercaptoethanol was added. RNA was size selected using RNA-Clean XP beads (Beckman Coulter) and full-length polyadenylated RNA was reverse transcribed using Superscript II. Resulting cDNA was amplified using 10 cycles and the amplified product was used to construct a pool of uniquely indexed samples using the Nextera XT kit (Illumina). Finally, pooled libraries were sequenced by Illumina NextSeq1000 (paired end).

### Statistical analysis

All data reported are represented as mean ± standard deviation. All measurements were made using distinct biological replicates and experiments characterizing individual sebaceous glands included several technical replicates per biological replicate. Prior experience in the lab on the number of mice needed to reach statistical significance in addition to mouse availability was used to determine sample sizes. All data being used in statistical comparisons were verified as normal using the Shapiro-Wilk measure of normality, and thus statistical significance was determined by a two-sided Student’s *t* test. Correlation analyses in Figure 5 were performed a Pearson correlation. All statistical analyses were performed using the R statistical computing environment version 4.2 and RStudio version 2022.02.1. For LCM-isolated sebaceous gland transcriptional analyses, *p* values were adjusted using the Bonferroni-Hochberg method and differential expression was determined as a gene with BH-adj *p* value < 0.05 and log_2_-fold change > 1 or < −1. For more exploratory analyses of global gene and pathway changes across generations in skin, small intestine, and liver, a less stringent cutoff was used of a non-adjusted *p* value < 0.05 and log_2_-fold change > 1 or < −1. For gametic small RNA and embryonic gene expression analyses, we also used a less stringent cutoff of a non-adjusted *p* value < 0.05 and log_2_-fold change > 1 or < −1. Correlation plot in Fig. 5H did not use a log_2_-fold change cut-off, to best show correlation of expression across all significant genes.

## Data and materials availability

Upon acceptance, the RNA sequencing datasets generated and analyzed in this study will be publicly available for download in the NCBI Gene Expression Omnibus (GEO). All data are available in the main text or the supplementary materials. The publicly available RNA sequencing dataset analyzed in Supplemental Figure 1A is available in the NCBI GEO at accession number GSE162925. Code is available upon request.

## Acknowledgments

**Non-author contributions**: We would like to thank members of the T.K. lab (M. Okumura), E.A.G. lab (A. Uberoi), and talented rotation students (M. Nelson, S. Barnett-Dubensky, and J. Doherty) for their assistance with carrying out experiments. We thank members of all the University of Pennsylvania and Children’s Hospital of Pennsylvania core facilities used including the Gnotobiotic Core (D. Kobuley and M. Albright), CHOP High-Throughput Sequencing Core (T. Orendovici and S. Mahoney), Penn Vector Core, and Skin Biology and Diseases Resouce-based Center, specifically the Cutaneous Phenomics and Transcriptomics core (S. Prouty and T. Dentchev). We also thank the Gnotobiotic Core from the University of North Carolina for coordinating the delivery of germ-free mice. For providing an open-source RNA sequencing analysis pipeline, we would like to thank the DIY Transcriptomics course (D. Beiting). Models created with BioRender.com.

## Author Contributions

Conceptualization: JCH, NAT, EAG, CCC, TK

Experimental design: JCH, NAT, EAG, CCC, TK

Hypothesis generation: JCH, NAT, EAG, CCC, TK

Methodology: JCH, NAT, BG, YY, LD, EKW, AH, CCC

Data analysis and interpretation: JCH, NAT, YY, EKW, CAT, EAG, CCC, TK

Funding acquisition: JCH, EAG, CCC, TK

Project supervision: EAG, CCC, TK

Writing – original draft: JCH, TK

Writing – review and editing: JCH, NAT, BG, LD, EKW, CAT, EAG, CCC, TK

## Extended Data

Figs. S1 to S4

**fig. S1.**
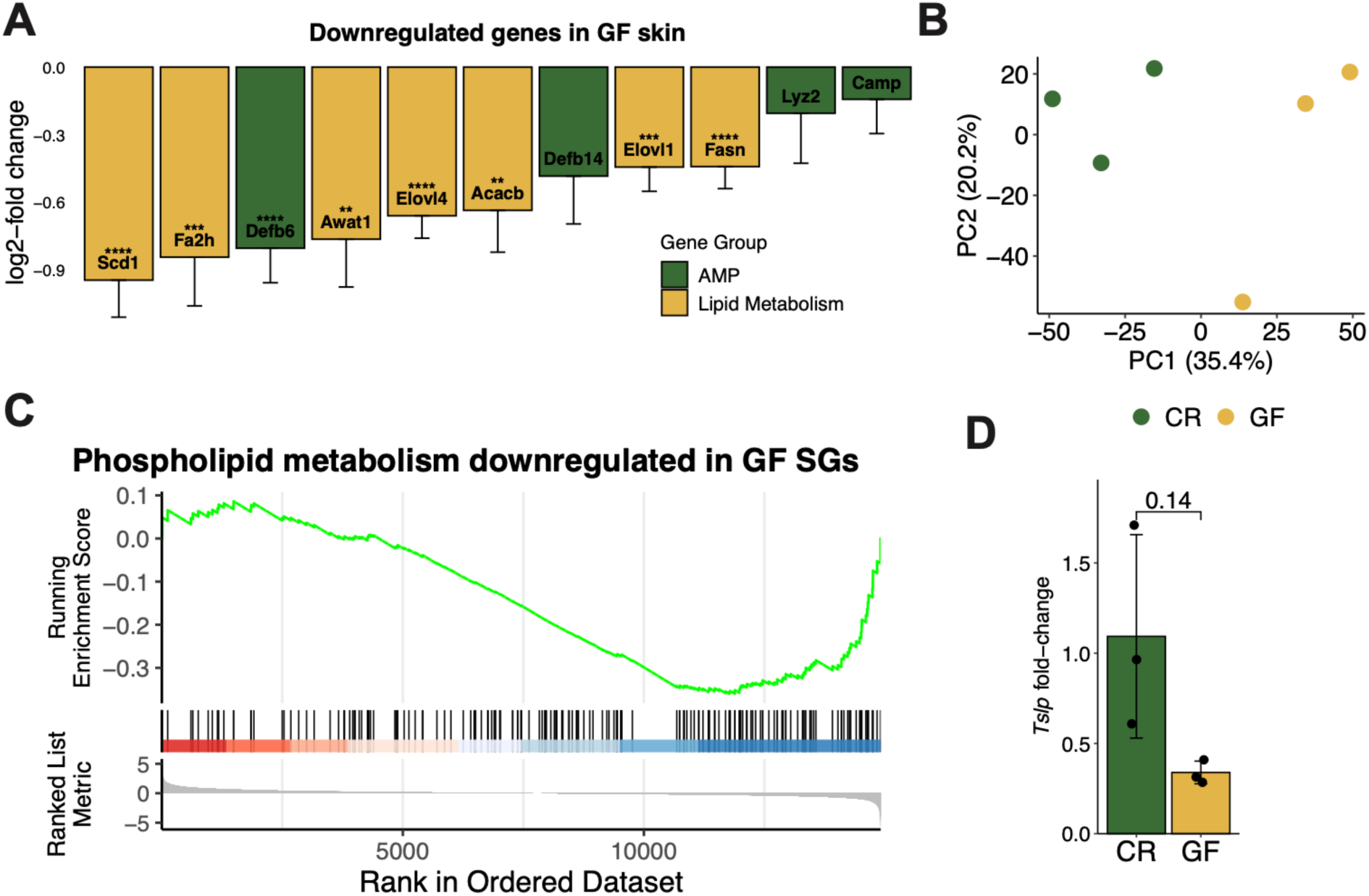
Skin and SG transcriptional profiles are dysregulated in GF mice. (**A**) mRNA expression of lipid-associated and antimicrobial peptide genes in GF mice compared to CR mice (*n* = 8 mice/group). *p* values are indicated on the y-axis, based on Fisher’s exact test and FDR-adjusted. (**B**) Principal component analysis showing principal component 1 (PC1) and PC2 for RNA-seq from the SG of CR or GF mouse skin (*n* = 3 mice/group). (**C**) GSEA plot displaying downregulated phospholipid pathway in the GF SG using BH-adjusted *P* value < 0.05. Genes shown in ranked order by running enrichment scores. (**D**) Skin mRNA expression of *Tslp* (*n* = 3 mice/group, qPCR normalized to *Gapdh* expression). No label, not significant, \*\**p* < 0.01, \*\*\**p* < 0.001, \*\*\*\**p* < 0.0001, by Fisher’s exact test, FDR-adjusted. Data are shown as mean ± SD.

**Fig S2.**
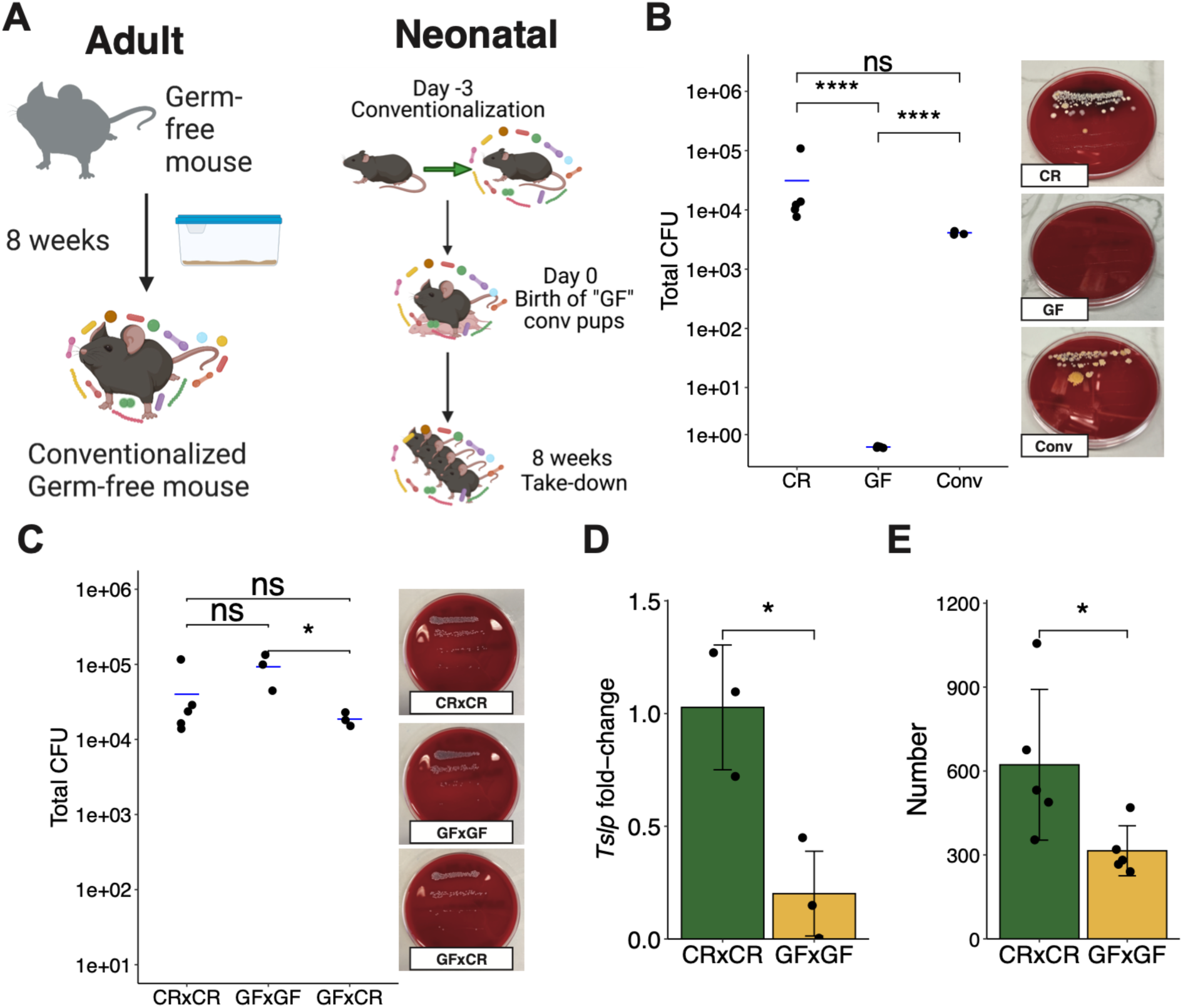
A similar bacterial burden in GF breeding combinations led to differential *Tslp* expression and T cell number in GF×GF F_1_ skin. (**A**) Schematic representing adult and neonatal germ-free conventionalization techniques. (**B**) Colony forming unit (CFU) quantification from back swabs of CR, GF and conventionalized adult mice post 8 weeks of colonization. (**C**) CFU quantification from back swabs of F_1_ CR×CR, GF×GF, and GF×CR mice. (**D**) F_1_ skin mRNA expression of *Tslp* (*n* = 3 mice/group, qPCR normalized to *Gapdh* expression). (**E**) Number of skin T cells in F_1_ mice by flow cytometry (*n* = 3 mice/group). All experiments performed 2-3 times. ns, not significant, \**p* < 0.05, \*\**p* < 0.01, \*\*\**p* < 0.001 by Student *t* test. Data are shown as mean ± SD.

**fig. S3.**
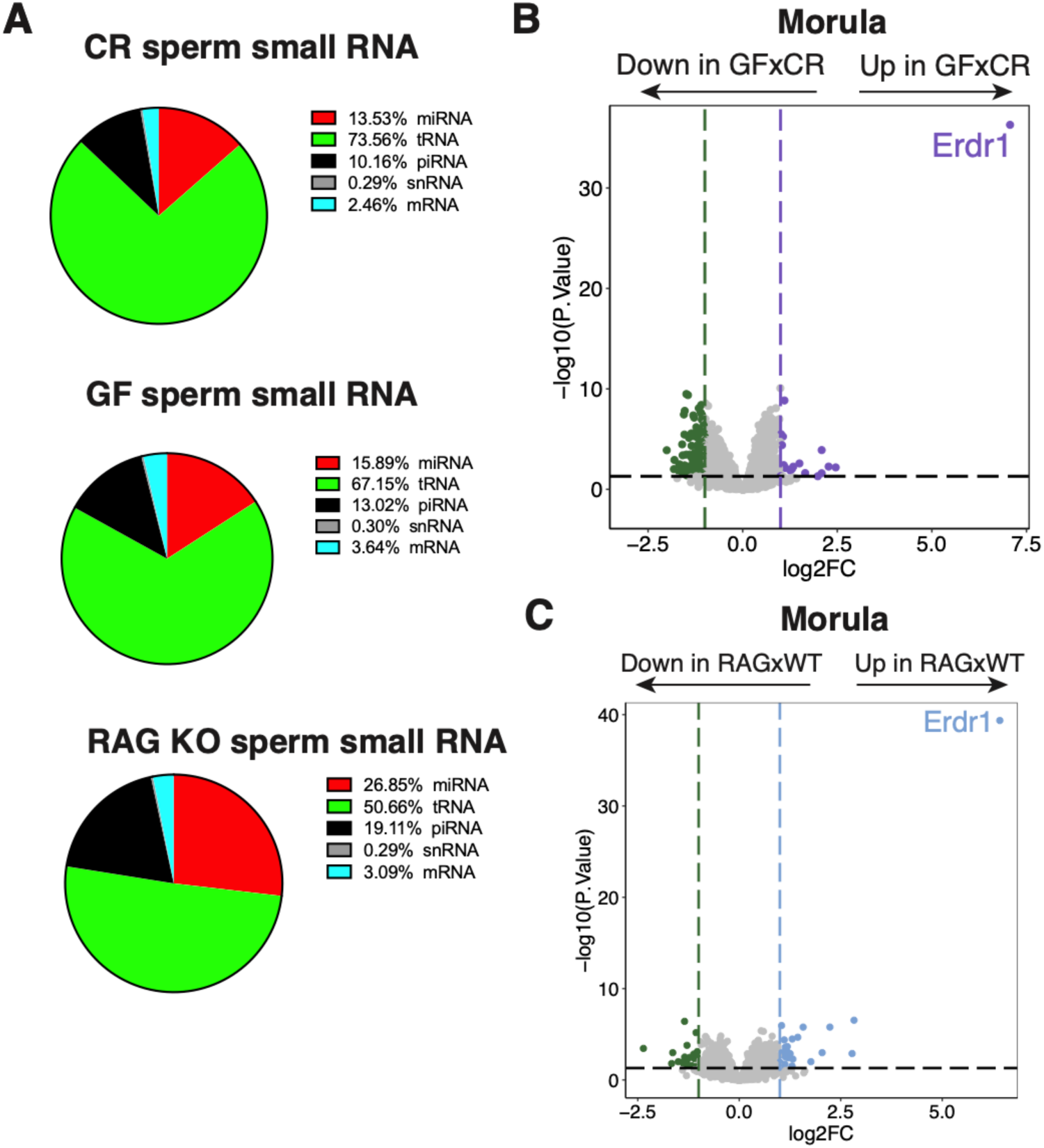
The microbiome and T cells can modulate levels of sperm small RNA and regulate gene expression in embryos. (**A**) Percentages of specific small RNAs present in CR, GF, and *Rag2^−/−^* sperm. (**B**) Gene expression by RNA-seq of CR×CR and GF×CR morulae (*n* = at least 17 morulae/group, collected over three biological replicates of IVF). (**C**) Gene expression by RNA-seq of WT×WT and RAG×WT morulae (*n* = at least 28 morulae/group, collected over three biological replicates of IVF). Sequencing experiments performed once.

**fig. S4.**
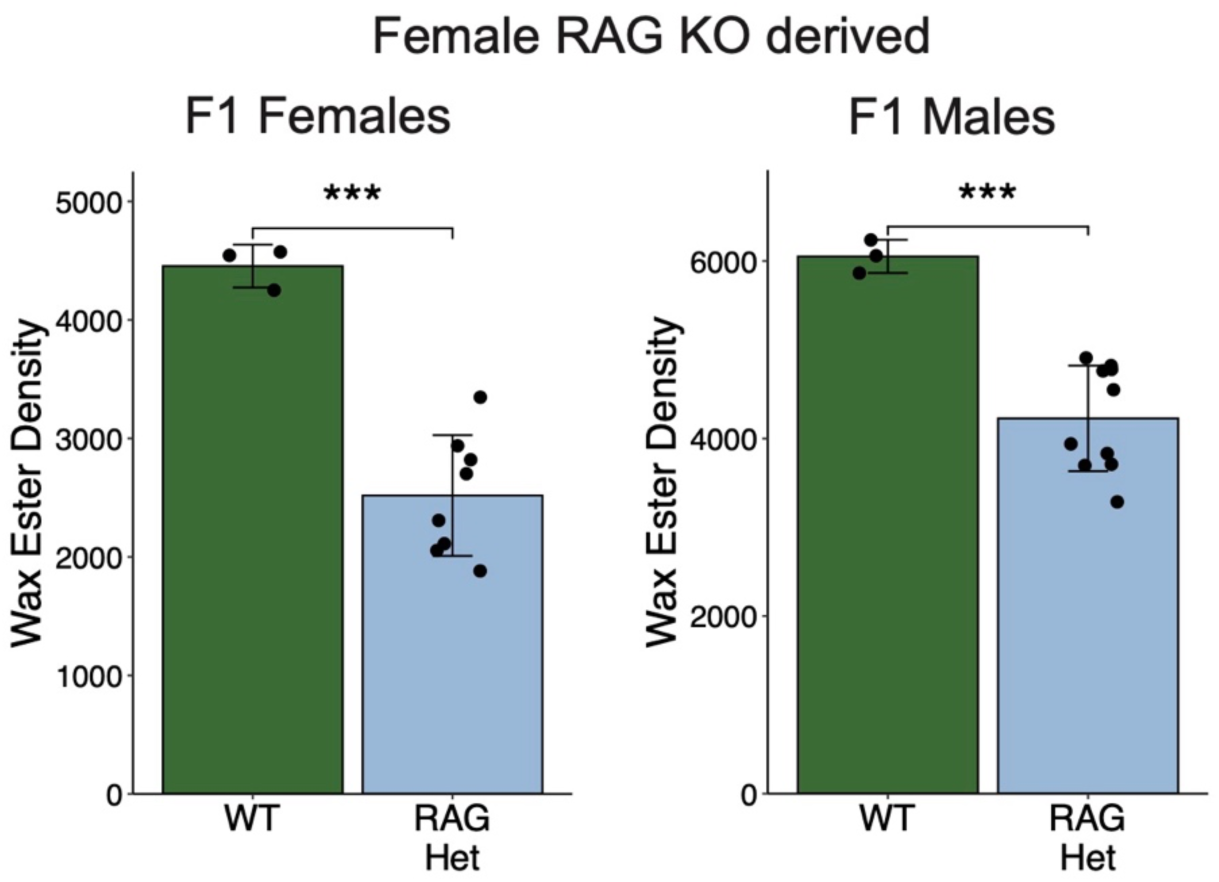
Progeny of *Rag2^−/−^* female mice crossed with WT male mice display a sebum secretion defect. TLC quantification of hair wax esters from female and male WT and F_1_ *Rag2^+/−^* mice (*n* = 3 (WT) or 8-10 (*Rag2^+/−^*) mice/group). Experiment performed twice. \*\*\**p* < 0.001 by Student *t* test. Data are shown as mean ± SD.

